# Intragenomic Conflict over Bet-Hedging

**DOI:** 10.1101/364190

**Authors:** Jon F Wilkins, Tanmoy Bhattacharya

## Abstract

Genomic imprinting, where an allele’s expression pattern depends on its parental origin, is thought to result primarily from an intragenomic evolutionary conflict. Imprinted genes are widely expressed in the brain and have been linked to various phenotypes, including behaviors related to risk tolerance. In this paper, we analyze a model of evolutionary bet-hedging in a system with imprinted gene expression. Previous analyses of bet-hedging have shown that natural selection may favor alleles and traits that reduce reproductive variance, even at the expense of reducing mean reproductive success, with the trade-off between mean and variance depending on the population size. In species where the sexes have different reproductive variances, this bet-hedging trade-off differs between maternally and paternally inherited alleles. Where males have the higher reproductive variance, alleles are more strongly selected to reduce variance when paternally inherited than when maternally inherited. We connect this result to phenotypes connected with specific imprinted genes, including delay discounting and social dominance. The empirical patterns are consistent with paternally expressed imprinted genes promoting risk-averse behaviors that reduce reproductive variance. Conversely, maternally expressed imprinted genes promote risk-tolerant, variance-increasing behaviors. We indicate how future research might further test the hypotheses suggested by our analysis.

## Introduction

Genomic imprinting is the phenomenon where the expression pattern of an allele depends on its parental origin (1). In the simplest cases, alleles at an imprinted locus are monoallelically expressed in a parent of origin–specific manner. This differential expression is the result of patterns of epigenetic modification that are established in the male and female germ lines, inherited by offspring, and propagated through development.

Although a number of explanations for the evolutionary origins of genomic imprinting have been proposed, the taxonomic and functional distribution of imprinted genes has best been explained by the Kinship Theory of Imprinting (2, 3), which attributes imprinting to an intragenomic conflict between maternally and paternally inherited alleles over the organismal phenotype. The Kinship Theory describes the conflict in terms of inclusive fitness effects, where an allele affects the fitness of multiple individuals in the population, some of which carry identical copies of the allele due to recent common ancestry. Because these probabilities of identity differ between maternally and paternally inherited alleles, there is a conflict between their inclusive fitness interests—an example of the “origin conflict” (4) category of intragenomic conflict.

The Kinship Theory has been particularly successful in providing a framework for understanding the effects of imprinted genes expressed in fetal and placental tissues during mammalian pregnancy. Gene expression in these tissues can influence the demand on maternal resources, affecting the fitness of the individual offspring as well as the number and overall fitness of the mother’s other offspring (including littermates in multiparous species as well as future offspring). The possibility of multiple paternity over the course of a female’s reproductive life means that the inclusive-fitness consequences of this resource demand impact maternally and paternally inherited alleles differently.

As a result, selection favors alleles that place greater demand on maternal resources when paternally inherited. This theory explains, in broad terms, the pattern of epigenetic silencing observed at imprinted loci with gene products that affect fetal and placental growth. Alleles at imprinted loci where increased expression leads to greater growth (via greater demand on maternal resources) are expressed only when paternally inherited, whereas alleles at growth-restricting loci are expressed when maternally inherited.

Imprinted gene expression is also found in adult tissues, particularly the central nervous system (5), where individual genes have been associated with a variety of cognitive and behavioral phenotypic effects (6, 7, 8). These include maternal care for offspring (9, 10), reactivity to novel environments (11), social dominance (12, 13), and delay discounting (14, 15) in animal models. Consequences of uniparental disomies and parental origin-specific deletions also point to an important role for imprinted genes in shaping cognitive and behavioral phenotypes in humans (16, 17, 18, 19, 7). While some interesting patterns are starting to emerge in terms of the types of cognitive and behavioral phenotypes that are affected by imprinted genes, the selective pressure(s) favoring the association of imprinted genes with these phenotypes have not yet been adequately identified.

### Bet-Hedging

Natural selection favors not only traits that increase mean reproductive success, but also traits that reduce reproductive variance, a phenomenon known as “bet-hedging” (20). Reproductive variance can arise from a number of different sources, such as spatial or temporal fluctuations in environmental conditions. Even in the absence of environmental variation, finite population sizes create a degree of reproductive variance. Gillespie (21, 22, 23) studied the effect of stochastic variation in individual reproductive success on the fate of alleles in the population. In Gillespies’s haploid, single-locus, two-allele model, the number of offspring produced by an individual carrying the *A*_1_ allele is drawn from a distribution with mean *μ*_1_ and variance 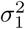. Similarly, the reproductive success of an *A*_2_ individual has mean *μ*_1_ and variance 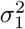. Allele *A*_1_ is expected to increase in frequency in the population if

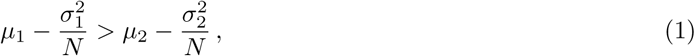

where *N* is the population size. This suggests that an allele with reduced mean fitness can invade the population, so long as it also provides a sufficiently large reduction in variance. It also implies that there is a continuum of allelic strategies that are characterized by different mean fitnesses, yet are selectively neutral with respect to each other. The trade-off between increased mean and reduced variance is determined by the population size *N*. The smaller the population size, the more important reproductive variance is to long-term evolutionary outcomes.

Frank and Slatkin (24) developed a more general framework that includes individual stochastic variation of the sort considered by Gillespie, as well as environmental fluctuations that lead to correlated changes in the reproductive success of individuals. For a two-allele system, they find that the expected change per generation in *q*_1_, the frequency of the *A*_1_ allele, *i.e.*, its proportion in the population, is approximately

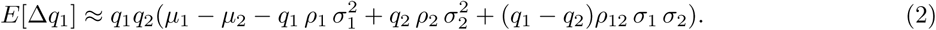

In this equation, *ρ*_1_ (*ρ*_2_) is the correlation of the reproductive success of two randomly chosen *A*_1_ (*A*_2_) individuals, and *ρ*_12_ is the cross-correlation between randomly chosen *A*_1_ and *A*_2_ individuals. Note that if individuals’ reproductive outputs are uncorrelated, *ρ*_1_ = 1*/*(*N*_*q*1_), *ρ*_2_ = 1*/*(*N*_*q*2_), and *ρ*_12_ = 0. The conditions under which *q*_1_ is expected to increase then reduce to those derived by Gillespie.

If the reproductive success of an allele is correlated across generations, these results need to be modified to include multigenerational effects. In this paper, we derive these modifications in the particular setting where dynamics over two generations are correlated. We also generalize the model to allow alleles to occupy different states (*e.g.*, male and female) for which the mean and variance of reproductive success may differ. Finally, we combine these two generalizations, allowing us to consider alleles at an imprinted locus, where the functional properties of an allele (and therefore its associated reproductive mean and variance) may differ, depending on the allele’s parental origin.

This analysis identifies a novel basis of intragenomic conflict over the trade-off between reproductive mean and variance. This conflict may have helped to shape the evolution of patterns of imprinted gene expression in the brain as well as the capacity for imprinted genes to influence specific cognitive and behavioral phenotypes. Following the presentation of the model, we discuss how these results relate to known behavioral effects of imprinted genes and how future research might test the model’s predictions.

### The Model

We present a brief recapitulation of the derivation of equation (2) in Supplementary Section S.1, both to clarify the assumptions that go into that derivation and to make subsequent derivations easier to follow. We denote the mean reproductive success of alleles *A*_1_ and *A*_2_, and of the population as a whole as

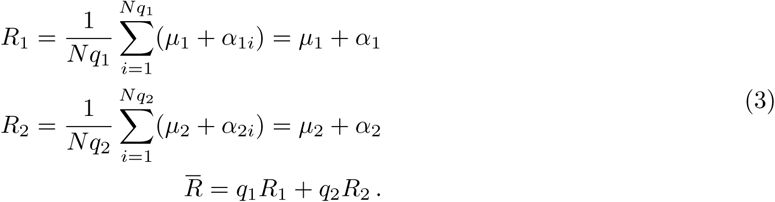

Here, *μ*_*k*_ are the expected mean reproduction for allele *k, α*_*ki*_ are the deviation of the reproductive success of *i*^th^ individual carrying allele *k* from this mean. In an infinite population of independently reproducing individuals, one expects *α*_*k*_ ≡ Mean[*α*_*ki*_] = 0, but in any finite realization, this mean deviation of the realized reproductive success from the expectation for allele *k* will typically be nonzero.

Without loss of generality, Frank and Slatkin set the mean reproductive success of the population, *q*_1_*μ*_1_ + *q*_2_*μ*_2_ = 1, which corresponds to rescaling all the *μ* and *α* by the expected reproductive success of the population. Equation (S2) and the assumption that the overall stochastic fluctuation in reproductive output is small relative to the total population size (Δ*N* ≪ *N*, or equivalently, *q*_1_*α*_1_ + *q*_2_*α*_2_ *≪* 1), leads to

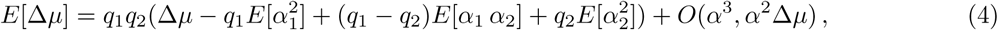

where Δ*μ* = *μ*_1_ − *μ*_2_. The details of the derivation are reproduced in Supplementary Section S.1. The derivation of equation (2) is completed when we substitute the definitions of the variances and covariances of the reproductive success provided in equation (S5). The result originally obtained by Gillespie arises as a special case if we assume that the reproductive success of different individuals is uncorrelated, in which case, the correlations arise purely from picking the same individual in a random sample of two individuals, and

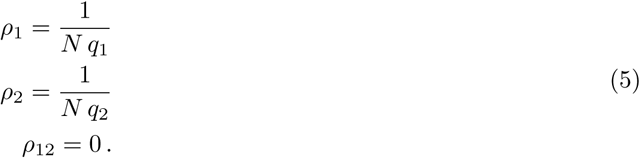

Substituting these values into equation (2) gives

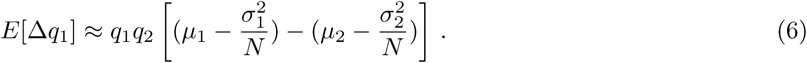

Note that if we think of the effective fitness of an allele as *w*_*e*_ = *μ - σ*^2^*/N*, equation (6) is simply the standard expression for the expected change in allele frequency under a fixed fitness difference. This effective-fitness definition also suggests the potential utility of the bet-hedging effective population size *N*_*bh*_, indicating the relative contributions of reproductive mean *μ* and variance *σ*^2^ to the long-term evolutionary fate of an allele under stochastic variation. In the haploid model described above, *N*_*bh*_ = *N*. In general, smaller values of *N*_*bh*_ would indicate that natural selection would more strongly favor traits that reduce reproductive variance.

### Diploidy, Imprinting, and Sex Differences

The first extension of this model is to a diploid system. For this and subsequent extensions, we assume fair segregation in heterozygotes. The diploid analog to equation (S4) is (see derivation of supplementary equation (S13) for details)

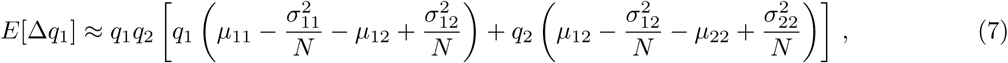

where *μ*_*ij*_ ≡ *μ*_*ji*_ and 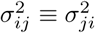 denote the scaled mean and variance of the reproductive fitness of an individual carrying the alleles *A*_*i*_ and *A*_*j*_. Again, using an effective fitness substitution 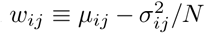, we can see that this equation reduces to

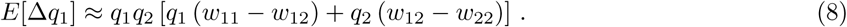

Note that this invokes a normalization step analogous to the one used to generate equation (S3). That is, the *μ* and *σ* terms are rescaled such that the mean reproductive output for the population is 1. In the diploid case, it is important to keep in mind that this quantity does not correspond to the expected number of offspring, rather it corresponds to the expected number of copies of the alleles passed on to the next generation. Due to diploidy, *μ* is, therefore, one half the expected number of offspring, and *σ*^2^, one fourth the variance in the number of offspring. These quantities correspond more closely to the mean and variance in the number of copies of a given allele passed on by individuals of the given genotype.

We can modify our diploid result to accommodate genomic imprinting by allowing the two heterozygous genotypes to have different phenotypes. From this point forward, we will use the convention that the subscripts are ordered such that the first number indicates the allelic identity of the maternally inherited allele, while the second indicates the identity of the paternally inherited allele. For example, *μ*_12_ denotes the mean reproductive success of an individual inheriting the *A*_1_ allele maternally and the *A*_2_ allele paternally.

The inclusion of imprinting leads to a minor modification of equation (7). Using the effective fitness notation, we have (see derivation of supplementary equation (S16) for details):

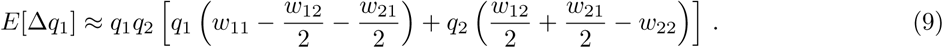

The only change is to replace the reproductive mean and variance of the heterozygous genotype with the average of the reproductive means and variances of the two phenotypically distinct heterozygotes.

At the simplest imprinted loci, one of the two alleles is transcriptionally silenced (in a parental origin-specific manner), and only the active allele affects the phenotype. In such a case, we can simplify equation (9). For example, for a locus where only the maternally inherited allele is expressed, *w*_11_ = *w*_12_ = *w*_1*_, and we have

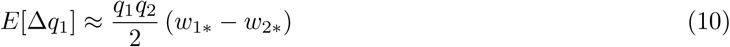

This is identical to what we get in the haploid case, but with the expected change in allele frequency reduced by a factor of two. This can be understood intuitively if we consider that one half of the alleles in the population have no effect on the phenotype, meaning that the efficacy of selection decreases by a factor of two.

The final extension we consider is to sex-specific phenotypes. The result is again straightforward. In the case of a 1:1 sex ratio (see derivation and discussion around supplementary equation (S20) for the more general case), the expected change in allele frequency is identical to the formulas presented above, with the effective mean and variance for the reproductive success given in terms of the means and variances in the two sexes by

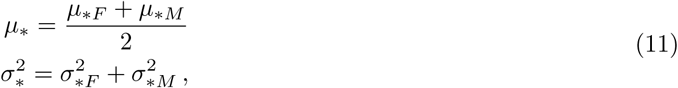

where the subscripts *F* and *M* represent the female and male phenotypes, respectively. The mean reproductive success of a genotype is the average of the mean reproductive success in males and females, while the reproductive variance is the sum of the male and female variances.

While we briefly treat the more general case in the supplementary information, we focus our analysis on the case where there are equal numbers of males and female for two reasons. First, we are particularly interested in applying our analysis to genomic imprinting in mammals, where population-wide sex ratios tend to be close to 1:1. Second, it is possible to assume equal numbers of males and females in the model without loss of generality, albeit at a cost to the interpretability of the model’s parameters. From the point of view of the model, an individual that leaves no offspring is no different from an individual that is never born. Therefore, we can equalize the sex ratio by introducing a number of non-reproducing “ghost” individuals of the less numerous sex. The null reproduction of these ghosts would then be taken into account when calculating reproductive mean and variance for that sex. The drawback of that approach is that these values measured in field studies need to be non-trivially transformed using the sex ratio to obtain the input parameter values for the model.

We note that, for each of the three extensions to the model, there is no qualitative change in the analysis. The expected change in allele frequency can be represented in the standard form in equation (6), where the means and variances of the two alleles are given by the average values of those quantities across the various states (genotype, parental origin, and sex) in which the alleles are found.

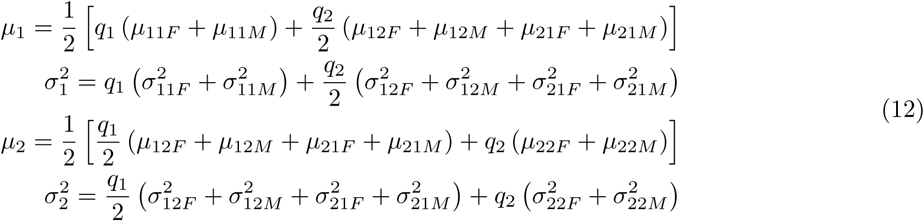

### Allele Frequency Change Across Two Generations

The overall reproductive success of alleles after two generations can be written in a form analogous to equations (3):

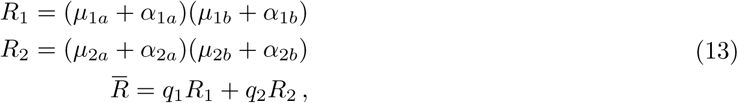

where the subscript *a* or *b* refers to the first or second generation, respectively. The frequency of allele *A*_1_ after two generations is then given by

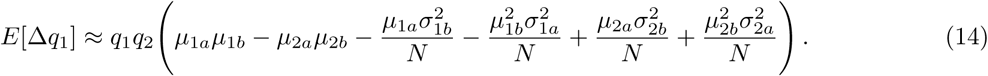

The details of the derivation are provided in Supplementary section S.4.

In considering the long-term evolutionary outcomes, it is useful invoke our a bet-hedging effective allele fitness. In the single-generation case, this was simply *w* = *μ - σ*^2^*/N*, and the analogous quantity for the two-generation case is

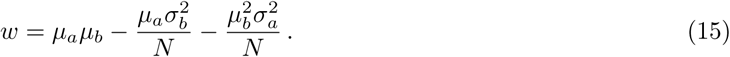

Note that in this expression, the variance in the second generation is multiplied by the mean of the first generation, while the variance of the first generation is multiplied by the square of the second-generation mean. This is easy to understand since the second generation reproductive success fluctuations of the first generation progeny are uncorrelated, leading to these second generation *variances* adding. Fluctuations in the first generation, however, are linearly inflated by the reproductive success in the second generation, so that the first generation variances getting multiplied by the *square* of the second generation reproductive success. A numerical simulation clearly shows the asymmetry between the two generations, and is shown in Fig. 1.

**Figure 1.**
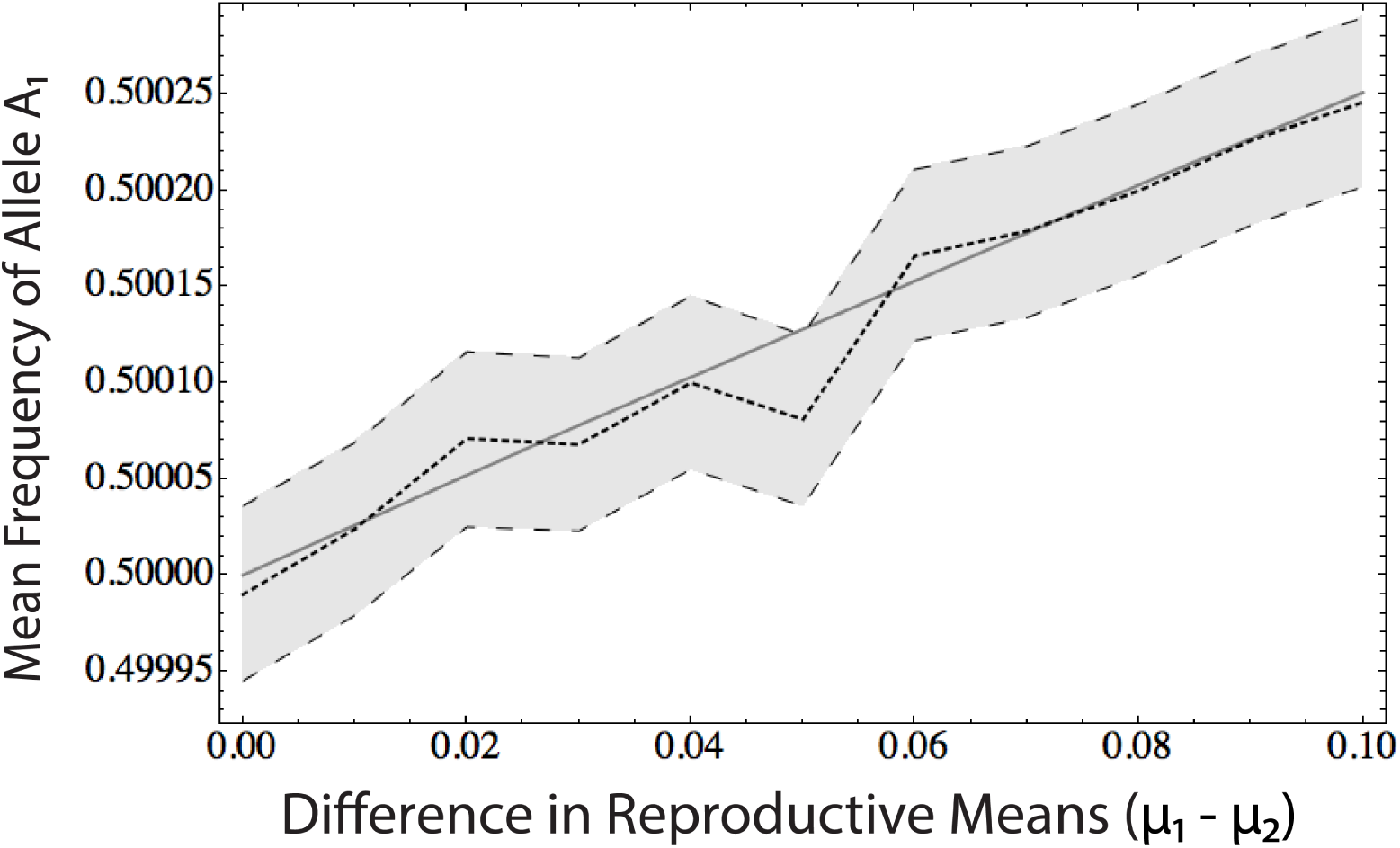
Two-generation fitness evaluated numerically. Two alleles are considered. Allele A_1_ has a mean reproductive success of *μ*_1_ and variance 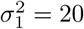 in the first generation, and mean *μ*_2_ and variance 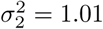 in the second. Allele A_2_ is like Allele A_1_, but with generations one and two interchanged. In all cases, (*μ*_1_ + *μ*_2_)*/*2 = 1. For various values of *μ*_1_ − *μ*_2_ plotted along the abscissa, 10,000,000 populations of size 1000, consisting initially of equal numbers of the two alleles, were simulated. The solid line is the expected frequency of A_1_ using the effective fitnesses from equation (15). The dark dashed line is the mean A_1_ allele frequency after two generations. The error bars are the 2-sigma confidence intervals on the mean.

### Two-Generation Bet-Hedging at an Imprinted Locus

Normally, we think of successive generations representing identical stochastic processes, and this is still true for the populations we consider. However, for an allele at an imprinted locus, there is a correlation between the states that an allele inhabits in subsequent generations. Specifically, an allele will only be maternally inherited in the present generation if it was present in a female in the previous generation. Likewise, paternally inherited alleles are always inherited from a male. This leads to interesting consequences, as we now argue.

The logic of the analysis that follows, and presented with mathematical details in Supplementary section S.5, can be grasped intuitively from consideration of equation (15). If the *a* terms in equation (15) represent the values in males (averaged across parental origin and genotype), then the *b* terms will represent values for paternally inherited alleles (averaged across sex and genotype). Similarly, if *a* represents females, *b* will represent maternally inherited alleles. Due to the final term in equation (15), the benefits of increasing mean reproduction (e.g., at the expense of increased reproductive variance) decline as the reproductive variance in the previous generation increases.

In most species, males have a higher variance of reproductive success than females. That means that in considering the fitness trade-off between increased mean and reduced variance, alleles will receive a greater benefit from reducing reproductive variance when paternally inherited, while alleles will receive greater benefit from increasing the mean when maternally inherited. At an imprinted locus, where alleles exhibit two distinct strategies based on parental origin, natural selection will favor divergent strategies, leading to the type of intragenomic conflict and evolutionary arms race observed in other imprinted systems. At the margin, paternally expressed imprinted genes will favor phenotypic traits that reduce reproductive variance (at the cost of reduced mean reproduction), while maternally expressed imprinted genes will favor traits that increase mean reproduction (at the cost of increased reproductive variance).

We can make this intuitive analysis explicit by first defining the overall reproductive mean and variance for an allele conditional on its being present in males or females. For allele *A*_1_ in females,

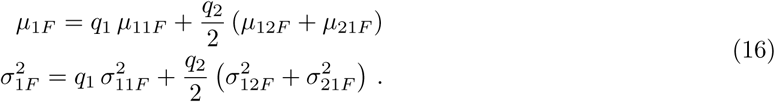

Analogous relationships hold for allele *A*_2_ and for males. We also have similar expressions for the reproductive mean and variance conditional on parental origin. Thus, for allele *A*_1_ when maternally inherited,

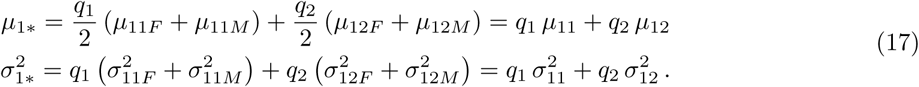

Again, the analogous expressions for *A*_2_ and for paternally inherited alleles are straightforward.

The two-generation effective fitness for allele *A*_1_ is simply the average of equation (15) over two sets of alleles. The first set comprises alleles that are present in females in generation *a* and are maternally inherited in generation *b*. The second set are of alleles that are present in males in generation *a* and are paternally inherited in generation *b*:

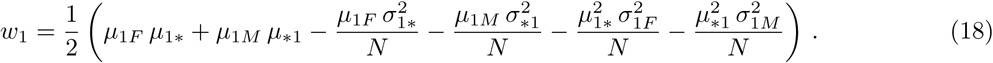

The term *w*_2_ can be similarly defined (see discussion following supplementary equation (S34) for details), and, with these definitions, the expected change in allele frequency is *E*[Δ*q*_1_] *≈ q*_1_*q*_2_(*w*_1_ − *w*_2_). In order to understand the basis of the intragenomic conflict, we compare this expectation for pairs of alleles *A*_1_ and *A*_2_ in two different contexts: an imprinted locus where only the maternally inherited allele is expressed *E*[Δ*q*_1_]*_m_*, and an imprinted locus where only the paternally inherited allele is expressed *E*[Δ*q*_1_]*_p_*.

For clarity of presentation, our analysis was restricted to the case where the alleles do not have sex-specific effects on mean reproductive success (*μ*_1*F*_ = *μ*_1*M*_ and *μ*_2*F*_ = *μ*_2*M*_), though it is easy to relax this assumption. The difference in the expected allele frequency changes is then given by

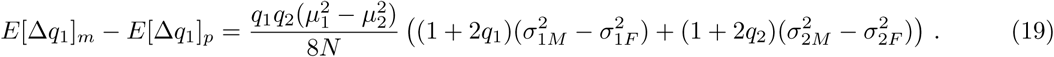

If reproductive variance is greater for males than for females, as is typically the case, then

*E*[Δ*q*_1_]*_m_ - E*[Δ*q*_1_]*_p_* will have the same sign as 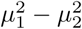. That is, if *μ*_1_ *> μ*_2_, allele *A*_1_ will have a greater advantage over allele *A*_2_ at a maternally expressed imprinted locus than at a paternally expressed one.

The result can perhaps be seen more clearly if we assume that 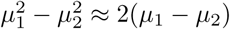, which follows if *μ*_1_ and *μ*_2_ are both close to 1, and we assume that the difference in male and female reproductive variances is the same for both alleles. We then have

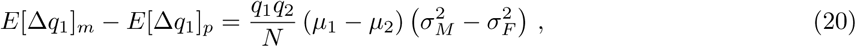

which is our final result discussed below.

## Results and Discussion

Population-genetic models of bet-hedging demonstrate that, all else being equal, natural selection will favor alleles that reduce individual reproductive variance. This establishes an evolutionary trade-off between reproductive mean and variance. An allele could have an evolutionary advantage, even if it leads to a reduction in mean reproductive success, so long as it also provides a sufficient reduction in reproductive variance. If we are talking about individual, stochastic variation in reproduction, the relative importance of mean versus variance is determined by the population size, with smaller population sizes corresponding to a greater evolutionary benefit to variance reduction.

The trade-off between mean and variance is illustrated in Figure 2. Natural selection will favor alleles that maximize *w*_*e*_ = *μ - σ*^2^*/N*. An evolutionarily stable strategy (ESS) will be one where the only evolutionarily accessible neighboring strategies would reduce *w*_*e*_. In the figure, that would be at the edge of the accessible strategies, where the slope *dμ/dσ*^2^ = *N*. The location of the ESS will depend on *N*, with a larger population size leading to higher values of *μ* and *σ*^2^ at equilibrium. The sensitivity of the location of the ESS to *N* will depend on the shape of the boundary.

**Figure 2.**
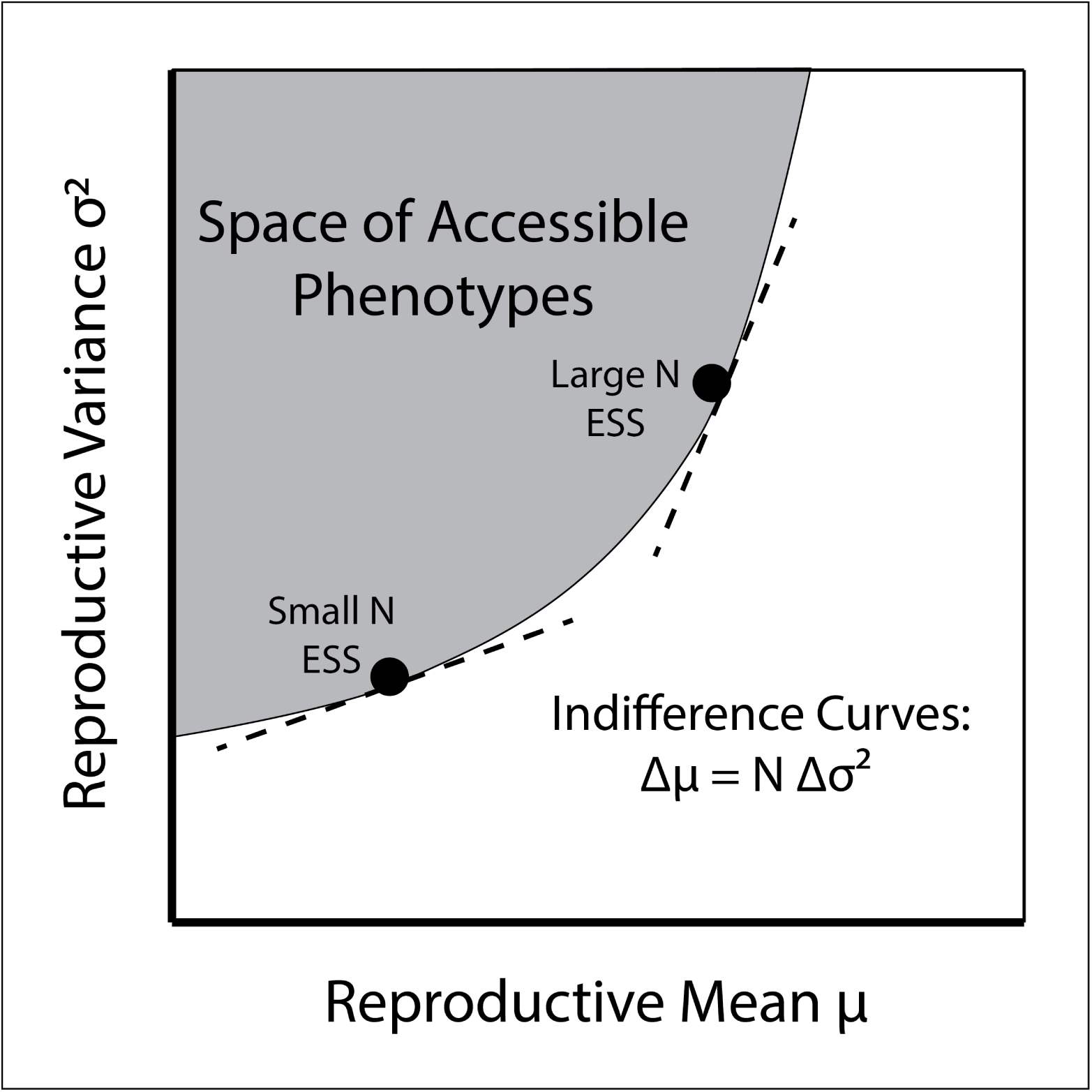
The bet-hedging trade-off between mean and variance leading to N-dependent Evolutionary Stable Strategies (ESS). This figure illustrates the trade-off between reproductive mean and variance. The plot represents the hypothetical space of combinations of reproductive mean and variance. Natural selection will favor traits that increase the mean *μ* (shift the phenotype to the right in the figure) and traits that reduce the variance *σ*^2^ (shift the phenotype down in the figure). Selection will be neutral with respect to traits that increase the mean and variance (or decrease the mean and variance) in a proportion set by the population size: when Δ*μ* = *N* Δ*σ*^2^. These indifference curves are indicated for small and large *N* by dashed lines. The shaded area represents the combinations of *μ* and *σ*^2^ that are evolutionarily accessible, while the unshaded area represents phenotypes that are inaccessible due to developmental, energetic, environmental, or other constraints. The evolutionarily stable strategy in such a system will be a point on the boundary of the set of accessible states where the slope of the boundary is *N*, as this is the point in the figure that maximizes the effective fitness *w*_*e*_ = *μ - σ*^2^*/N*.

The analysis developed here demonstrates that the inclusion of imprinted gene expression potentially disrupts this ESS. Equation (S32) shows that, if there is a difference between the variances in reproductive success between the two sexes, the evolutionary trade-off between mean and variance differs between maternally and paternally inherited alleles. If the reproductive variance is greater in males than in females, as is typically the case, then mean reproductive success will be marginally more important for maternally inherited alleles, while reproductive variance will be marginally more important for paternally inherited alleles. We can see this by imagining a pair of alleles *A*_1_ and *A*_2_ for which *E*[Δ*q*_1_]*_p_* = 0, such that these alleles would be selectively neutral at a paternally expressed imprinted locus. If the same pair of alleles were instead at a maternally expressed imprinted locus, natural selection would favor whichever of the two alleles had the greater value of *μ*, despite the fact that this allele would also have a higher value of *σ*^2^.

Thus, with genomic imprinting, we no longer have a purely stable ESS. Assuming the boundary of accessible strategies is smooth, if we start at the ESS for an unimprinted locus, the population will be subject to invasion by alleles at imprinted loci. If 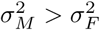, there will exist accessible strategies that increase *μ* and *σ*^2^ that will be favored by natural selection on alleles at a maternally expressed locus. Likewise, there will be accessible strategies that reduce *μ* and *σ*^2^ that will be favored by natural selection acting on a paternally expressed locus.

Figure 3 illustrates how imprinting alters the conceptual model presented in Figure 2. Sex differences in reproductive variance lead to differences in the bet-hedging effective population size. Higher reproductive variance in males (females) leads to a reduced bet-hedging effective population size for paternally (maternally) inherited alleles. This sets the stage for an arms race, with maternally expressed imprinted genes accumulating effects that increase reproductive mean and variance and paternally expressed imprinted genes accumulating effects that reduce reproductive mean and variance.

**Figure 3.**
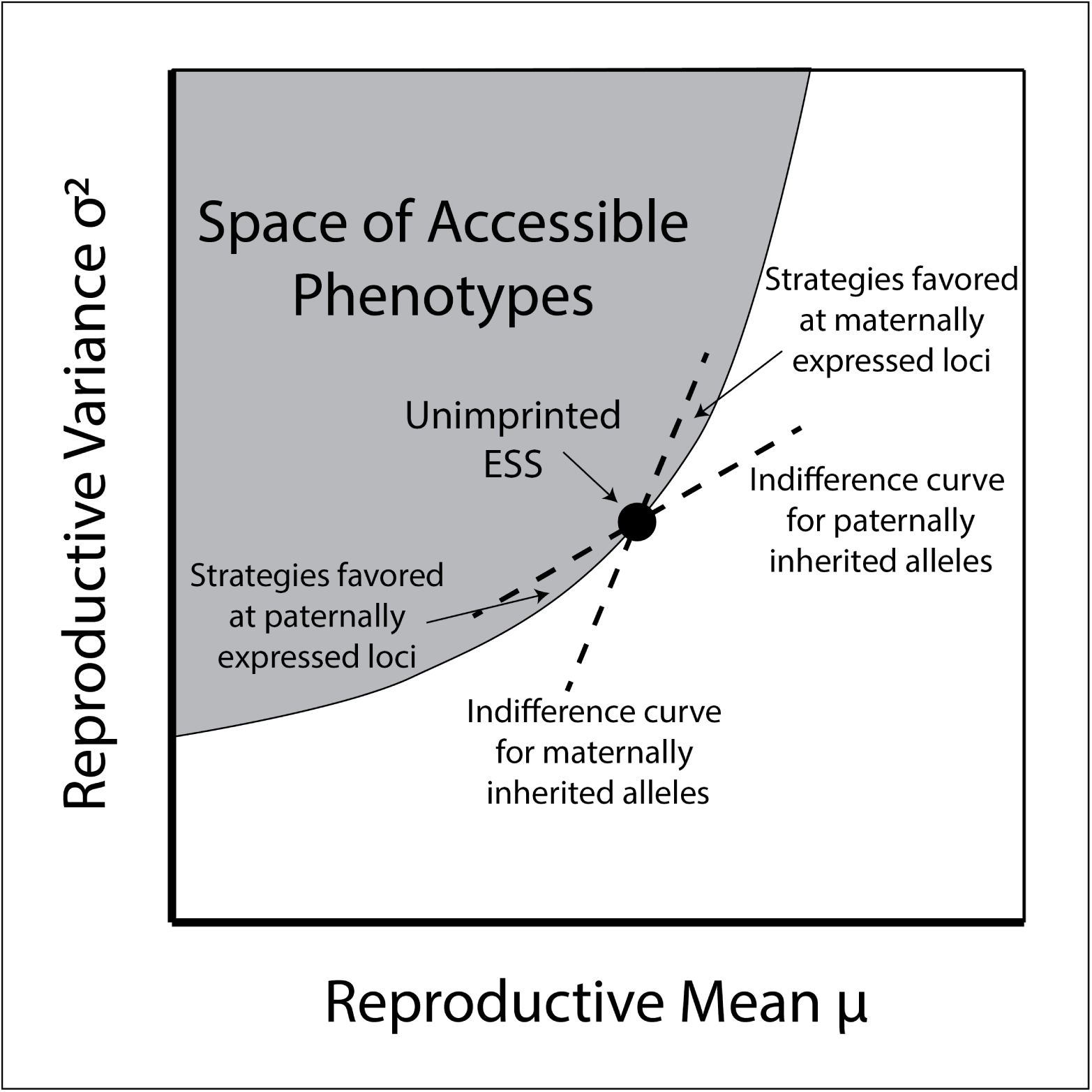
Intragenomic conflict over bet-hedging. This figure illustrates how the conceptual model from Figure 1 changes in the presence of imprinted gene expression in a system where males have a higher variance of reproductive success than females. Because maternally and paternally inherited alleles have different bet-hedging effective population sizes, the slope of the corresponding indifferences differ. Between the two indifference curves are strategies that would be favored by alleles at a maternally expressed locus, but not a paternally expressed one (or vice versa). Under higher male reproductive variance, the strategies favored only at maternally expressed loci are associated with increased reproductive mean and variance, while strategies favored only at paternally expressed loci are associated with reduced reproductive mean and variance.

### First-Order Consequences of Second-Order Effects

An important caveat to this analysis is that the selection asymmetry identified here is of the order of the selection difference between alleles (*μ*_1_ − *μ*_2_) divided by the population size *N*. The rule of thumb in population genetics is that the efficacy of selection depends on the product of the strength of selection *s* associated with a new allele and the population size *N*. If *Ns* < 1, the allele is effectively neutral, because the stochastic influence of genetic drift outweighs the systematic differences in reproduction due to selection. From this perspective, it may seem reasonable to suspect that the selection effects described here will be too modest to have any real explanatory power.

However, we note that this systems is somewhat different from the contexts in which this rule of thumb is typically invoked. Typically, one assumes a Poisson-distributed number of offspring, giving rise to the variance in the number of offspring being related to the mean reproductive success, *σ*^2^ *≈ μ*. In such a situation, a *O*(1*/N*) difference in *μ* is of the same order of magnitude as the *σ*^2^*/N* term in the effective fitness, and one can not analyze selective advantage by considering the mean dynamics alone. In this analysis, we are explicitly considering the stochastic dynamics, and our results are thus properly sensitive to these possibly small differences. Note also that we are not speaking here about selection acting in opposition to mutational entropy (where there are many more accessible mutations acting in opposition to the direction of phenotypic change favored by selection). Rather, we start from a circumstance in which there is a continuum of very nearly neutral phenotypes in the form of a fitness ridge defined by the indifference curve set by the population size *N*. We might expect alleles to be constrained to this ridge, but following a random walk along it.

The phenomenon described here might best be understood as introducing small, opposing biases into this random walk at maternally and paternally expressed imprinted loci. Even with a small bias, we should expect, over sufficient time, to see the accumulation of substantial opposing phenotypic effects. This accumulation is the natural outcome of intragenomic arms races, resulting from the fact that the effects of maternally and paternally expressed alleles manifest through a shared organismal phenotype. Each time a mutation fixes at a maternally expressed locus that increases reproductive mean and variance, it increases the opportunity for selection to favor a mutation at a paternally expressed locus that reduces them. Similarly, fixation of favored allele at a paternally expressed locus increases the opportunity for selection to act at a maternally expressed one.while such an arms race will lead to the accumulation of opposing phenotypic effects at maternally and paternally expressed imprinted loci (25), it may have only a modest effect on the wild-type reproductive mean and variance, a question we do not consider here. Evidence of the escalation will primarily be in the form of the large phenotypic perturbations associated with loss-of-function mutations at imprinted loci affecting the trait. Specifically, the prediction is that loss-of-function mutations at maternally expressed imprinted loci would result in a phenotypic shift in a direction associated with reduced reproductive variance, while loss-of-function mutations at paternally expressed loci would be associated with increased reproductive variance.

Note that the interpretation of such loss-of-function mutations is not always trivial, as the evolutionary model is focused on changes of small phenotypic effect. In the wake of an evolutionary arms race, loss-of-function mutations may result in a large perturbations associated with phenotypic “overshoot” (26). The phenotypic consequences of major genetic perturbations should therefore be treated with caution when inferring the evolutionary history of the system.

While this coevolutionary escalation may have little effect on the population mean phenotype (including the distribution of reproductive success, risk preference profiles, etc.), the precise nature of the evolutionary consequences will depend on the details of how various genes interact to construct that phenotype. Depending on the genes’ pleiotropic effects, evolutionary equilibrium could be characterized by the fixation of maladaptive traits, including traits that are the object of the intragenomic conflict as well as pleiotropically linked traits over which maternally and paternally inherited alleles share the same optimal phenotype (27). Escalation may also lead to phenotypic decanalization, resulting in the persistence of extreme phenotypes at high frequency in the population (28).

### Delay Discounting and other Risk-Related Traits

The model developed here applies to the lifetime mean and variance of reproductive success. However, there is value in identifying short-term cognitive and behavioral traits that might be expected to correlate with these quantities and that are amenable to experimental manipulation. This expected correlation involves the use of the term “risk” to bridge two distinct, but related, concepts. The first is behavioral risk, where a particular benefit to the organism is associated with a potential cost. This cost could come in the form of death or injury, or simply in the form of an opportunity or energetic cost. The second is “risk” in the sense of reproductive variance. Implicit in the interpretation of our model’s results that follow is the assumption that there is a causal relationship between behavioral risk and reproductive risk. For example, a behavior that involves an increased risk of premature death will increase the probability that that individual produces no additional offspring. A behavior that has the possibility of increasing or decreasing an individual’s social status may increase or decrease it’s future reproductive output, thereby increasing that individual’s reproductive variance.

In general, we expect increased reproductive mean and variance to be positively correlated with the pursuit of high-risk, high-reward behavioral strategies. In that case, maternally expressed imprinted genes are expected to favor these high-risk strategies, while paternally expressed imprinted genes will favor risk-averse behaviors that lead to more modest, but more certain, rewards.

Delay discounting is one model for probing behavioral risk preferences. In a typical experiment, an individual makes a choice between receiving a small reward that is delivered immediately and a larger reward that is delivered with some delay. As that delay increases, the individual becomes less and less likely to choose the larger reward. The connection to risk follows from the assumption that, in the natural environment, a delayed reward is a less-certain reward. Waiting for a delayed reward increases the possibility that the reward will be lost (*e.g.*, to a competitor), or that the individual will be placed in danger (*e.g.*, due to predation). The probability of choosing the larger reward, as a function of the time delay, defines the individual’s “delay-discount function.”

Within the delay-discounting context, the model predicts that maternally expressed alleles should favor greater willingness to wait for a larger reward, even if that waiting imposes risk. That is, these alleles will favor a shallower, or less-steep, discount function. Conversely, paternally expressed alleles will favor a steeper discount function, corresponding to less willingness to tolerate a delay in order to secure the larger reward.

The effects of two imprinted genes, *Nesp* and *Grb10*, on delay discounting in mice have found effects that are consistent with these predictions (see Figure 4). *Nesp* encodes a maternally expressed transcript within the complex imprinted *Gnas* locus (29) and is expressed primarily in the hypothalamus and midbrain (11). Dent et al. (14) characterized the delay-discounting behavior in mice whose maternally inherited *Nesp* allele had been inactivated by a targeted genetic deletion. The result was that these mice were less willing to wait for a larger, delayed reward compared with controls. In the risk interpretation of delay discounting, the loss of maternally expressed *Nesp* resulted in more risk-averse behavior, consistent with the model’s predictions that maternally inherited alleles would be selected to favor greater risk tolerance.

**Figure 4.**
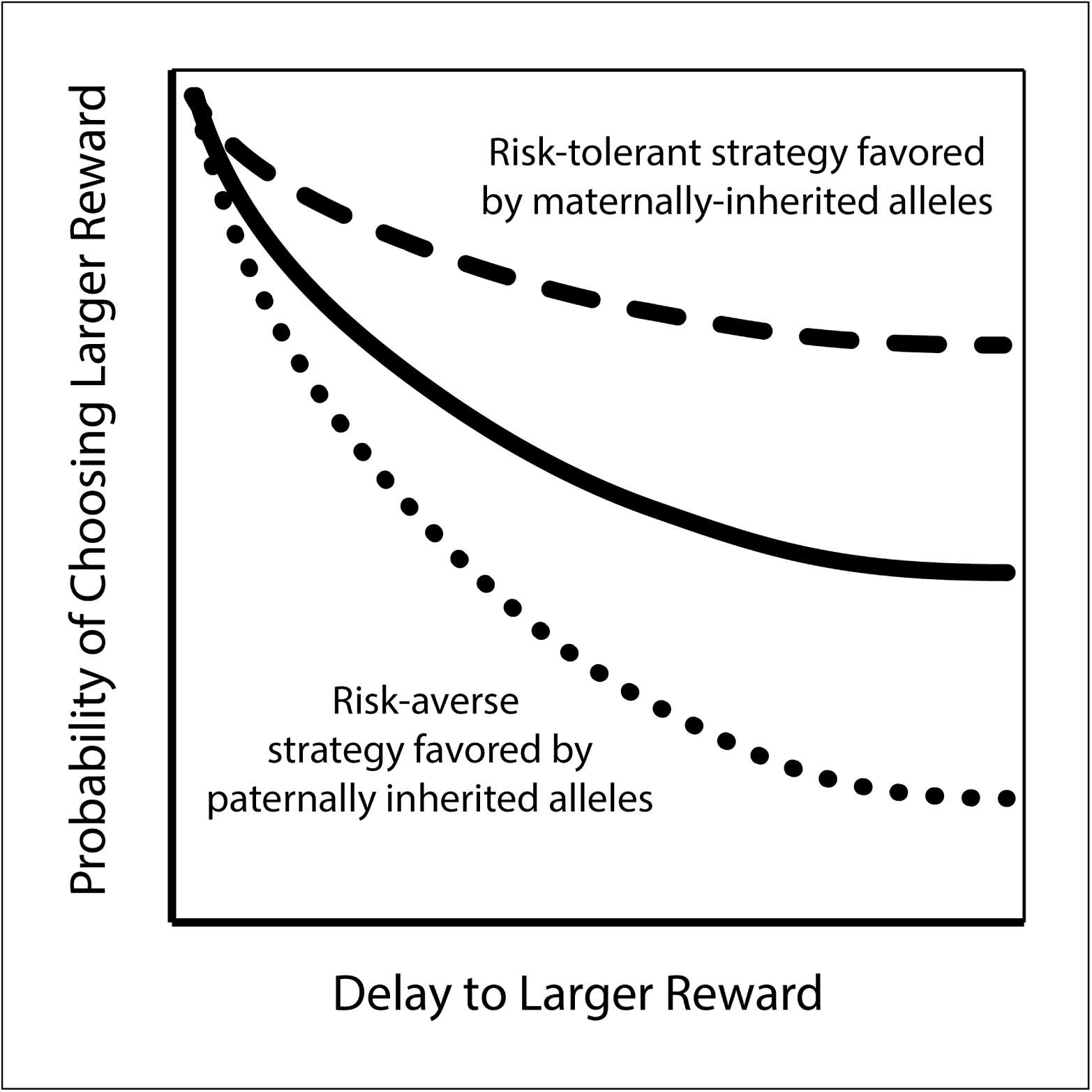
Intragenomic conflict over delay discounting. This figure schematically illustrates the effect of imprinted genes on delay discounting behavior in mice. The solid line represents wild-type discounting behavior. Mice choose between a small reward that is received immediately and a larger reward that is received after some delay. The longer the delay, the less likely the larger, delayed reward will be chosen. Elimination of the paternally expressed *Grb10* shifts behavior to the dashed line, indicating a greater willingness to tolerate delay (and risk) in favor of the larger reward. Elimination of the maternally expressed *Nesp* shifts behavior to the dotted line, producing a risk- (and delay-) averse phenotype.

In an analogous set of experiments, Dent et al. (15) analyzed the delay-discounting behavior in mice whose paternal expression of *Grb10* had been eliminated in the brain. *Grb10* is maternally expressed during development, and in peripheral tissues in adults, but is expressed exclusively from the paternally inherited allele in the central nervous system (30, 12). Elimination of expression from the paternally inherited allele was found to increase willingness to wait for a larger, delayed reward. This result is consistent with *Grb10* having been selected to favor more risk-averse behaviors when paternally inherited.

*Nesp* and *Grb10* have also been associated with other risk-related behaviors *Nesp* has also been associated with reactivity to novel environments, with maternal null mice showing increased activity when placed in a novel environment, but a reduced willingness to explore when given a choice between novel and familiar environments (11). This is potentially consistent with a role for maternally expressed *Nesp* in promoting risk-tolerant behaviors, in this case a willingness to explore an unfamiliar environment.

Elimination of paternal expression of *Grb10* results in a specific increase in social dominance behavior (12), as does an increase in the expression of the maternally expressed *Cdkn1c* gene in the brain (13). Assertion of social dominance can be interpreted as a high-risk behavioral strategy, as it may lead to increased reproductive success (if the animal achieves social dominance), or may lead to negative reproductive consequences as a result of social conflict. This is consistent with natural selection acting in opposite directions in these oppositely imprinted genes. Paternally expressed *Grb10* in the brain has been selected to reduce risky, variance-increasing social dominance behavior, whereas maternally expressed *Cdkn1c* has been selected to enhance social dominance behavior.

Two paternally expressed imprinted genes, *Peg1/Mest* and *Peg3*, have been associated with maternal behavior towards offspring (9, 10). In both cases, loss of paternal expression of the gene in the brains of adult females leads to deficits in specific maternal-care behaviors, including nest building and pup retrieval. One interpretation of maternal care behavior is that it modulates a trade-off between number and quality of offspring. A reduction in maternal care would allow the mother to focus her resources on producing her next litter, at the expense of increasing the probability that pups in her current litter do not survive, thereby potentially increasing both the mean and variance of her reproductive output. That would imply that the normal function of these paternally expressed genes in the female adult brain is to reduce reproductive variance by focusing resources on ensuring the survival of a smaller number of offspring.

### Other Evolutionary Explanations of Behavioral Effects of Imprinted Genes

Several proposals for the basis of the evolution of imprinted gene expression in the brain have been put forward. In general, the models are consistent with at least a subset of the empirical data. The coadaptation theory of imprinting proposes a synergistic matching of phenotypes between offspring and mothers, including, potentially, maternal behaviors (31). Sex-biased dispersal will create relatedness asymmetries within local social groups, which can then generate asymmetric selection on incest avoidance (32), maternal care (33), and on dispersal behavior itself (34, 35). More generally, demographic and social structures will lead to intragenomic conflict over helping behaviors, including maternal care, alloparenting, and generalized altruism (36, 37, 38).

Determining which proposal, or combination of proposals, is best able to account for the cognitive and behavioral effects of imprinted genes is difficult at the moment. Behavioral phenotypes are complex, sensitive to context, and shaped by interactions among various genetic and environmental factors that are not fully understood. The fitness effects associated with specific behaviors may also depend on complex social and environmental patterns that may or may not be adequately reproduced in the laboratory setting. Furthermore, most imprinted genes that are expressed in the brain are also expressed in early development, where they affect growth and development. The behavioral effects of simple genetic deletions are then confounded by major developmental perturbations. In fact, maternally expressed *Nesp* and paternally expressed *Grb10* are among the only imprinted transcripts for which the effects of standard genetic manipulations are limited to the central nervous system.

### Origins of Imprinted Genes Affecting Behavior

While there is widespread imprinted gene expression in the mammalian central nervous system, imprinted genes are best known for their effects on early growth and development, particularly during pregnancy. In the context of fetal growth, the selection asymmetry derives from two stylized facts: 1) the resources supporting early growth come preferentially from the mother, and 2) offspring of the same mother may have different fathers. While the magnitudes of these underlying asymmetries may have varied over time, their polarity has likely been constant over the history of mammals. The result is a selection asymmetry on maternally and paternally inherited alleles that has been sufficiently strong and persistent to drive the evolution of the molecular machinery of imprinting and the acquisition of imprinted gene expression at dozens to hundreds of loci.

The strength of the selection asymmetry in fetal growth is likely due to the fact that relatively modest alterations to resource allocation early in development can have large effects on an offspring’s growth, survival, and eventual reproduction. By contrast, the selection asymmetry on gene expression in the central nervous system is likely much more subtle. While there is a relatively straightforward connection between, say, growth-factor expression in the placenta and the acquisition of maternal resources, gene expression in the adult brain has its fitness effects via complex behavioral, social, and demographic structures.

It seems unlikely, then, that selection asymmetries on maternally and paternally inherited alleles expressed in the central nervous system would be strong and persistent enough to drive the acquisition of imprinting at a previously unimprinted locus. Indeed, we find that the imprinted genes expressed in the brain are largely a subset of those expressed in fetal and placental tissues. The few exceptions (e.g., *Nesp* and paternal expression of *Grb10*) are transcripts produced from complex loci where most imprinted transcripts are expressed in early development.

This pattern is consistent with a “growth-first” model of imprinting evolution (39), where the relatively large intragenomic conflict associated with prenatal growth drives the acquisition of imprinted gene expression at individual loci. Once a locus has become imprinted, the evolution of its expression and the function of its gene product(s) are shaped by different selection pressures than the rest of the (unimprinted) loci in the genome. More subtle selection asymmetries may alter the timing, level, and location of expression in ways that differ for maternally and paternally expressed loci. These imprinted genes may gradually modify their functions and/or acquire new functions in a way that enhances the evolutionary success of the expressed allele.

It may be that, “Why have genes affecting traits like risk become imprinted in the brain?” is the wrong way to ask the question. These genes likely became imprinted because of their effects on fetal growth, and they may or may not have had their current behavioral effects at that time. The better question may be, “Why have these imprinted genes in the brain become associated systematic effects on risk (and other behaviors)?” This framing points to the value of comparative analyses across taxa. If we look in the brains of species without imprinting (or where imprinting is at least less prevalent), do these genes play analogous developmental roles? Are they expressed in the brain? If so, do they have similar effects on cognition and behavior? Answering these questions will allow us to better understand the evolutionary sequence leading to the current state.

### Empirical Predictions and Tests

Ultimately, the value of any evolutionary model is determined by its capacity for making sense of empirical patterns. The model developed here has the potential to make sense of imprinted gene effects on risk-related behaviors. There is an emerging pattern where alleles at maternally expressed imprinted loci favor greater risk tolerance, while paternally expressed loci favor risk aversion. This pattern is not yet as extensive or robust as, say, the pattern of fetal growth effects of imprinted genes. Further genetic and behavioral work will be required to determine the generality of the patterns in terms of both number of loci and taxonomic breadth.

An advantage of this model is that it depends on a pattern that is widespread and that has likely been common over the necessary evolutionary timescale: the asymmetry between male and female reproductive variances. Due to the subtlety of the selection asymmetry, the underlying variance asymmetry needs to have existed consistently for a long time in order for the arms race to have driven the accumulation of significant imprinted gene effects on the target phenotype. Among mammals, this variance asymmetry has likely been consistent and nearly universal.

Exceptions to the pattern of variance asymmetry provide opportunities for hypothesis testing. For example, the naked mole-rat, *Heterocephalus glaber*, exhibits a division of reproductive labor somewhat like that found in eusocial insects (40). Colonies consist of dozens to hundreds of individuals, but have only a single breeding female. Male reproduction is also skewed, but not to the same degree, with 1-3 breeding males per colony, and a more rapid turnover of breeding males than breeding females (41). The naked mole-rat is therefore a rare example of a mammal in which females likely have a higher reproductive variance than males. In this lineage, then, we would expect the direction of the selection asymmetry over risk-related behaviors to have been reversed.

Differences in the magnitude of the variance asymmetry between pairs of closely related species also provide opportunities for hypothesis testing. The expectation is that the larger the difference between male and female reproductive variance, the more intense the arms race. While a high-asymmetry species and a low-asymmetry species may have similar risk profiles, the two would be the result of different balances of underlying imprinted gene effects. A high-asymmetry species would have a balance between genes of large effect, while a low-asymmetry species would have a balance between genes of small effect.

If such pairs of species can be identified, the model developed here would make predictions regarding reciprocal heterosis in the risk preferences of hybrid offspring. For example, if a male from a high-asymmetry species were crossed with a female from a low-asymmetry species, we would expect the offspring to inherit paternally expressed alleles of large effect, and maternally expressed alleles of small effect. Because maternally expressed imprinted genes are predicted to promote risk tolerance, while paternally expressed imprinted genes will promote risk aversion, we should expect these hybrids to exhibit a risk-averse phenotype. The reciprocal cross (of a low-asymmetry female with a high-asymmetry male) would be expected to be more risk tolerant.

The power of the empirical tests described here depends not only on the identification and characterization of species conforming to specific demographic profiles, but also on the evolutionary lability of the genes and traits in question. As we noted early in the discussion, the selective pressure described here is weak, and we expect it to lead to the accumulation of measurable genetic and phenotypic traits only long evolutionary times. If the pattern of reproductive variance in a species changes, it may take considerable time for that change to be reflected in the function or expression of imprinted genes that affect risk. While the naked mole-rat may have a reproductive variance that is higher in females than in males, mole-rats in general (family Bathyergidae) exhibit a wide range of social structures (41). If the social structure of the naked mole-rat is a recently derived trait, selection may not yet have significantly altered the relevant imprinted genes. On the other hand, the diversity of social structures among mole-rats may provide interesting opportunities for reciprocal hybridization experiments.

## Conclusion

Our analysis identifies a novel basis for intragenomic conflict over the evolutionary trade-off between reproductive mean and variance. When males have a higher variance of reproductive success than females, as is commonly the case, alleles are more strongly selected to reduce reproductive variance when paternally inherited than maternally inherited. We have connected this result to empirical data on imprinted gene effects on risk-related behaviors. Consistent with the predictions of the model, there is a trend in which paternally expressed genes favor risk-averse behaviors, while maternally expressed genes favor more risk tolerance. Additional comparative and molecular data will be required in order to fully evaluate the extent to which this conflict can account for the taxonomic and phenotypic distribution of imprinted gene effects in the brain.

## Acknowledgments

We thank Anthony Isles, Andy Gardner, and the anonymous reviewers who provided feedback on various versions of the manuscript over time. The development of this work was supported in part by Leverhulme Trust Project Grant (F/00 407/BF).

## Supplementary Information

### S.1 The Model

We recapitulate briefly the derivation of equation (2) to explain the notation and to clarify the assumptions that go into that derivation. For ease of comparison, we follow the nomenclature of Frank and Slatkin. The mean reproductive success of alleles *A*_1_ and *A*_2_, and of the population as a whole can be written as

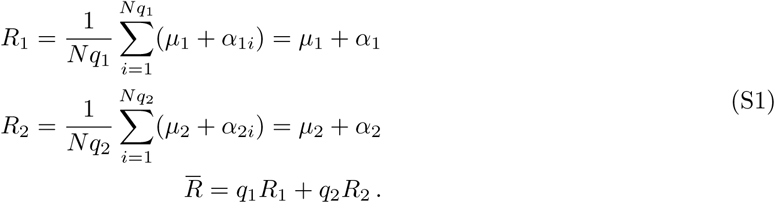

Here, *μ*_*k*_ are the expected mean reproduction for allele *k, α_ki_* are the deviation of the reproductive success of *i*^th^ individual carrying allele *k* from this mean, and *α*_*k*_ ≡ Mean[*α*_*ki*_] represent the mean deviation of the realized reproductive success from the expectation for allele *k*.

The frequency 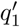 of the *A*_1_ allele in the next generation is given by 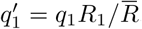. The expected change in the frequency of the *A*_1_ allele, is simply 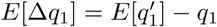. Substituting the relations from equations (S1) and using *q*_1_ + *q*_2_ = 1, we have

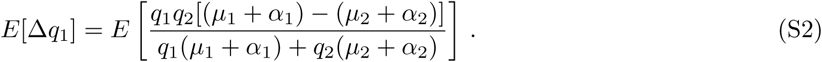

without loss of generality, Frank and Slatkin set *q*_1_*μ*_1_ + *q*_2_*μ*_2_ = 1, which corresponds to rescaling the *μ* and *α* by the expected reproductive success of the population. Equation (S2) then reduces to

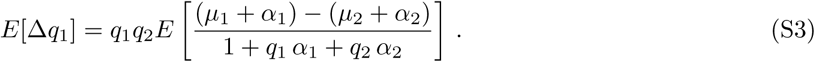

The trouble in evaluating equation (S3) comes from the fact that there is not a general solution for the expectation of a ratio. However, provided that the overall stochastic fluctuation in reproductive output is small relative to the total population size (Δ*N* ≪ *N*, or equivalently, *q*_1_*α*_1_ + *q*_2_*α*_2_ *≪* 1), we can approximate it as

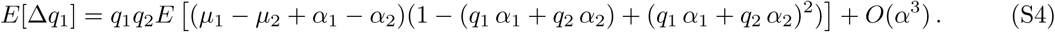

One potentially confusing aspect of this derivation arises from the fact that we are considering two different types of mean value. The first, represented by *α*_1_ and *α*_2_, is an average across individuals, but limited to a single realization of the stochastic process undergone by the population as a whole. The second, represented by the expectation operator *E*[*…*], is a mean taken over many hypothetical realizations of the reproductive process of the entire population. While *α*_1_ and *α*_2_ could each take on either positive or negative values in any single realization, these quantities are defined such that *E*[*α*_1_] = *E*[*α*_2_] = 0. The higher moments of these fluctuations are, however, not zero:

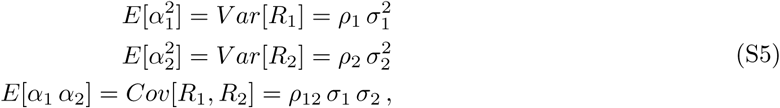

where *V ar*[*R*_*k*_] and *Cov*[*R*_1_, *R*_2_] represent the variance of *R*_*k*_ and the covariance of *R*_1_ with *R*_2_, respectively.with this in mind, we can now take the expectation of each of the terms in equation (S4). We can also simplify notation by introducing Δ*μ* = *μ*_1_ − *μ*_2_. This gives

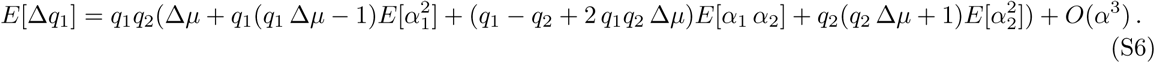

To arrive at their final expression, Frank and Slatkin make the further assumption that Δ*μ* is small. Equation (S4) then reduces to

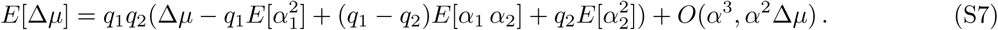

### S.2 Diploid Model

For the diploid analog of the Frank and Slatkin model, we begin with the reproductive success of the three genotypes. For simplicity, we assume a well mixed population at Hardy-Weinberg equilibrium.

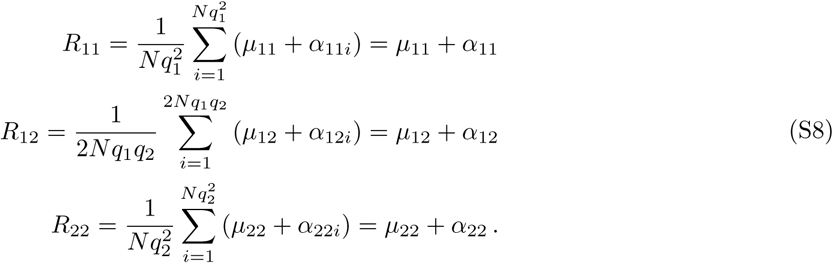

From this, we can calculate the reproductive success of the two alleles:

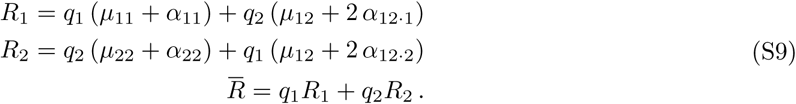

Note that in these expressions, we have split the *α*_12_ term into two components. In the previous expression, *α*_12_ represents the deviation of the number of offspring from the mean for *R*_12_ individuals. Assuming fair segregation, we the expected number of those offspring to carry the *A*_1_ an *A*_2_ alleles will be equal *α*_12·1_ represents the excess number of offspring to inherit an *A*_1_ allele from an *R*_12_ parent (and *α*_12·2_ the excess inheriting *A*_2_). The quantity *μ*_12_*/*2 + *α*_12·1_ is distributed as a binomial *B*(*μ*_12_ + *α*_12_, 1*/*2), and *α*12·2 = *α*_12_ − *α*_12·1_.

Now we can write our expression for the expected change in allele frequency

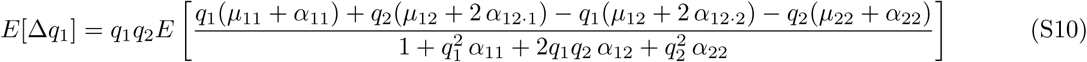

where we have again normalized by the total expected reproductive output by setting 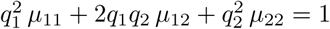. As in the haploid case, we assume that the fluctuation Δ*N* is small compared with the total population size *N*. This allows us to approximate this ratio as

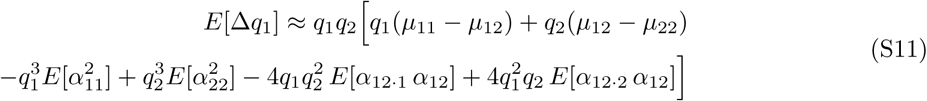

we again assume that the reproductive success of different individuals is uncorrelated, and we additionally assume that the population is at Hardy-Weinberg equilibrium. In this case, the non-zero variance and covariance terms are

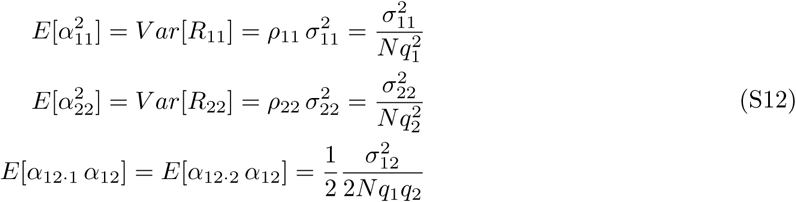

The approximation for the change in allele frequency then reduces to

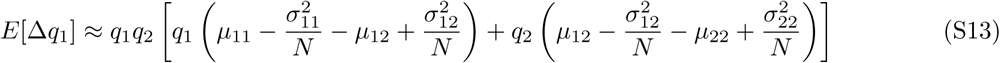

Note that, as in the haploid analysis, if we define the effective fitness of a genotype as *w*_*e*_ = *μ - σ*^2^*/N*, this reduces to the standard formula. Furthermore, if allelic effects on reproductive mean and variance are additive, it reduces to the haploid equation.

### S.3 Genomic Imprinting

Treatment of the general diploid case with genomic imprinting requires the addition of the fact that the two heterozogous genotypes may have different phenotypes. To accommodate this, we modify our notation such that two-number subscripts are ordered, with the first number indicating the identity of the maternally inherited allele and the second number indicating the identity of the paternally inherited allele. For example *μ*_12_ now represents the mean reproductive success of heterozogous individuals whose *A*_1_ allele was maternally inherited and whose *A*_2_ allele was paternally inherited.we now have four expressions for the mean reproductive output in a given generation:

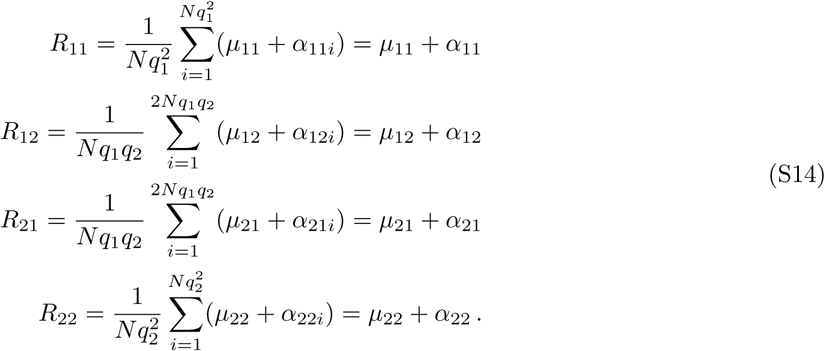

The corresponding expressions for the reproductive output of the two alleles are

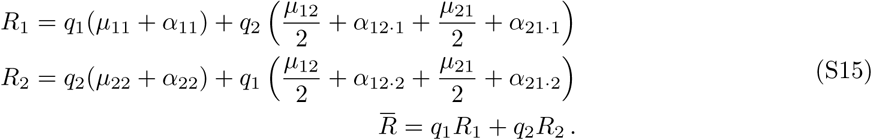

Once again, we assume that the fluctuations in total reproductive output are small compared with the population size, and that individual reproductive outputs are uncorrelated. The expected change in allele frequency then becomes

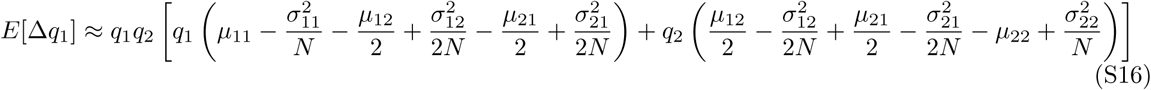

The result is virtually identical to what we found in the diploid case (without imprinting), with the mean and variance of the heterozygotes’ reproductive output being replaced by the averages of the means and variances of the two different heterozygotes. That is, we once again recover the standard expression for the change in allele frequency if we define effective fitnesses as 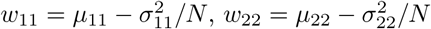, and 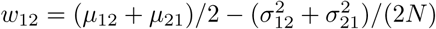.

Often, genomic imprinting involves the transcriptional silencing of one of the two alleles, such that the phenotype of the individual depends only on the maternally inherited (or paternally inherited) allele. For example, if we were considering an imprinted locus with expression only from the maternally inherited allele, we would only need to consider two values of reproductive mean and variance: 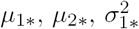, and 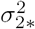, where * indicates either allele. Equation (S16) then reduces to

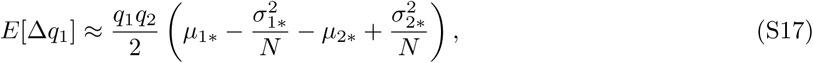

which is identical to the haploid result, except that the expected change in allele frequency is reduced by a factor of two.

### Two Sexes

We now consider the case where our diploid model has two sexes, in which the same genotype may be associated with different means and variances of reproductive success. To indicate the two sexes, we will include an additional subscript of *F* or *M* to each of the variables previously introduced. As before, we start with the mean reproductive success of each type in the population:

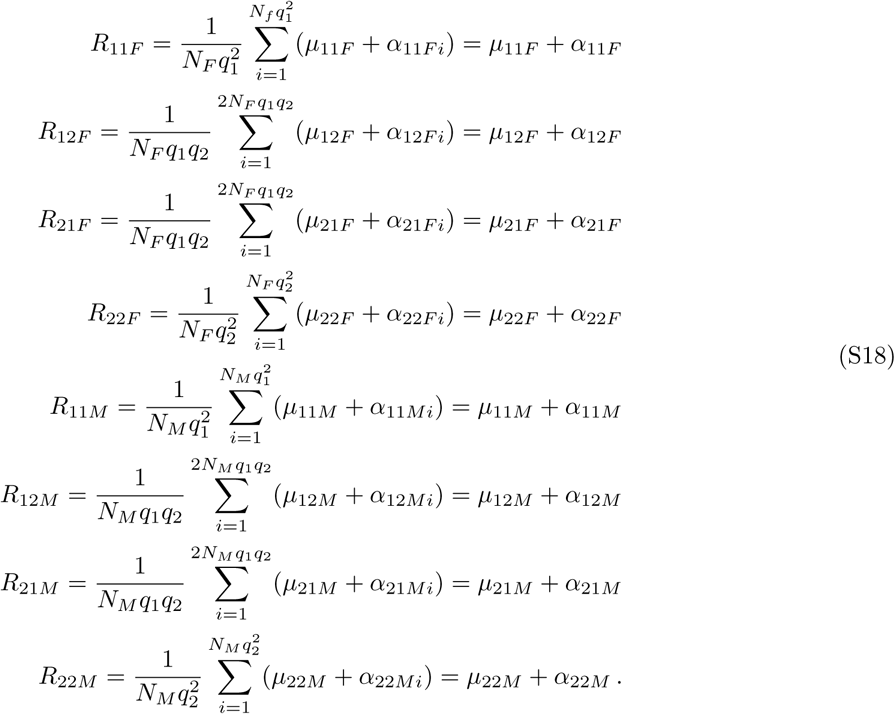

Because the total reproductive output of males and females in the population must be equal, the allele frequency in the next generation will simply be the average of the frequencies of the alleles passed on by males and females. Assuming that mating and sex determination are both independent of the genotype at this locus, this means that we can analyze the two sexes separately. That is,

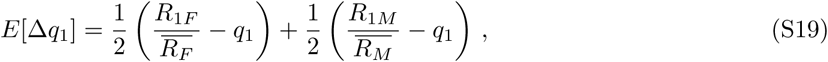

where the *E*[Δ*q*_1_]*_*_* terms have the same form as equation (S16). Substituting the appropriate expressions into this equation gives us

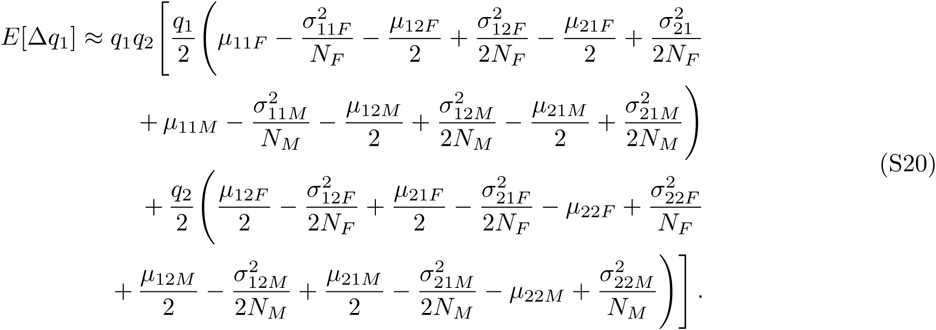

Recall that our earlier analysis involved normalizing the mean and variance by the total expected reproductive output 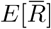. In this analysis, that normalization happened separately for males and females. That is, if we want to interpret *μ* and *σ*^2^ as the mean and variance of the number of offspring, we would need to make the following changes:

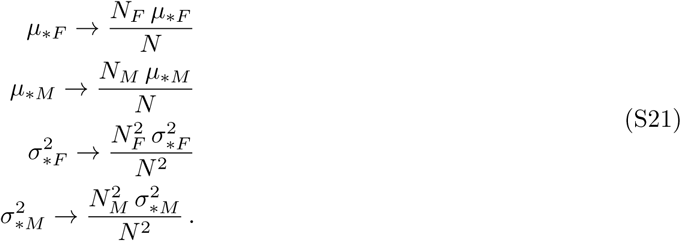

we will not make that substitution here, since the separately normalized versions of *μ* and *σ*^2^ correspond more closely with our intuitive notions of relative fitness.

Our analysis will focus on the case where the sex ratio is 1:1 (*N*_*F*_ = *N*_*M*_), however, we pause to note a few interesting features of equation (S20). First, the effective mean reproductive success of a genotype is simply the arithmetic mean of the genotype’s relative fitness in males and females: *μ*_*\**_ = (*μ*_*\*F*_ + *μ*_*\*M*_)*/*2. The effective reproductive variance of a genotype, by contrast, is a weighted average that depends more heavily on the reproductive variance in the rarer sex.

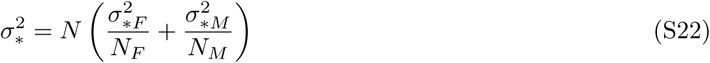

Note that this also means that selection will favor bet-hedging phenotypes more in the rarer sex. Given that the benefits of variance reduction are greater in smaller population sizes, this is not surprising. In the case of a 1:1 sex ratio, these effective mean and variance terms can be substituted to recover equation (S16).

### S.4 Two-generation model

The frequency of allele *A*_1_ after two generations follows from equation (13) and is given by

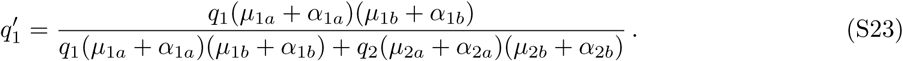

In order to simplify this expression, we introduce the following definitions:

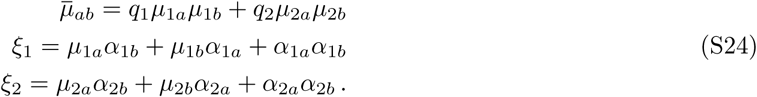

Substituting this notation into equation (S23) gives us an expression for the change in allele frequency after two generations:

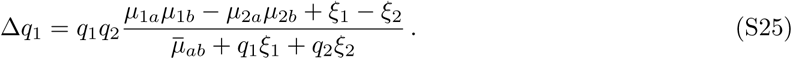

In this form, we can see that equation (S25) is analogous to equation (2). The expected reproductive success of alleles *A*_1_ and *A*_2_ is *μ*_1*a*_*μ*_1*b*_ and *μ*_1*a*_*μ*_1*b*_, respectively. The terms *ξ*_1_ and *ξ*_2_ are the deviations from those mean values. We scale *μ*_*ab*_ to one as before, and we assume that the difference between the means is small (*|μ*_1*a*_*μ*_1*b*_ − *μ*_2*a*_*μ*_2*b*_*| ≪* 1) and that the total stochastic deviation is small relative to the mean (*|q*_1_*ξ*_1_ + *q*_2_*ξ*_2_*| ≪* 1). We also restrict our analysis to the case where the reproductive success of different individuals is uncorrelated. The expected value of the change in allele frequency is then given approximately by

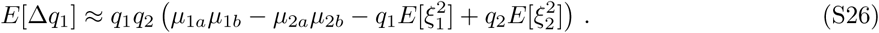

Substituting the relationships from equation (S24) back in and taking the expectations, as above, yields

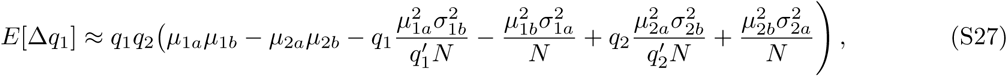

where the primes denote the allele frequencies in the intermediate generation, and we have discarded terms of order 1*/N* ^2^. Recognizing the fact that 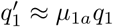 to the required order, this further reduces to

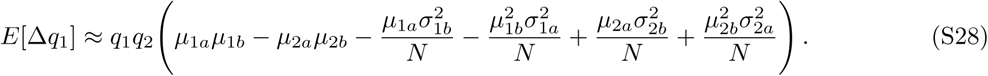

### Two-Generation Bet-Hedging at an Imprinted Locus

The logic of the analysis that follows can be grasped intuitively from consideration of equation (15). If the *a* terms in equation (15) represent the values in males (averaged across parental origin and genotype), then the *b* terms will represent values for paternally inherited alleles (averaged across sex and genotype). Similarly, if *a* represents females, *b* will represent maternally inherited alleles. Due to the final term in equation (15), the benefits of increasing mean reproduction (e.g., at the expense of increased reproductive variance) decline as the reproductive variance in the previous generation increases.

In most species, males have a higher variance of reproductive success than females. That means that in considering the fitness trade-off between increased mean and reduced variance, alleles will receive a greater benefit from reducing reproductive variance when paternally inherited, while alleles will receive greater benefit from increasing the mean when maternally inherited. At an imprinted locus, where alleles exhibit two distinct strategies based on parental origin, natural selection will favor divergent strategies, leading to the type of intragenomic conflict and evolutionary arms race observed in other imprinted systems. At the margin, paternally expressed imprinted genes will favor phenotypic traits that reduce reproductive variance (at the cost of reduced mean reproduction), while maternally expressed imprinted genes will favor traits that increase mean reproduction (at the cost of increased reproductive variance).we can make this intuitive analysis explicit by first defining the overall reproductive mean and variance for an allele conditional on its being present in males or females. For allele *A*_1_ in females,

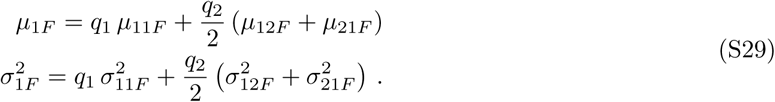

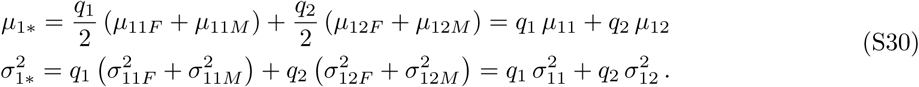

The two-generation effective fitness for allele *A*_1_ is simply the average of equation (15) over two sets of alleles. The first is alleles that are present in females in generation *a* and are maternally inherited in generation *b*. The second is alleles that are present in males in generation *a* and are paternally inherited in generation *b*:

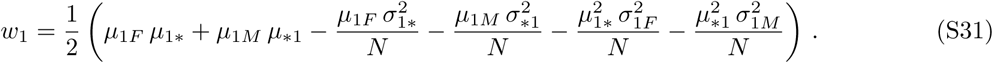

For clarity of presentation, our analysis is restricted to the case where the alleles do not have sex-specific effects on mean reproductive success (*μ*_1*F*_ = *μ*_1*M*_ and *μ*_2*F*_ = *μ*_2*M*_), though it is easy to relax this assumption. The difference in the expected allele frequency changes is then given by

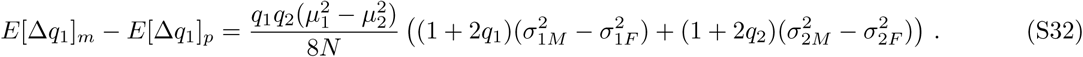

If reproductive variance is greater for males than for females, as is typically the case, then *E*[Δ*q*_1_]*_m_ - E*[Δ*q*_1_]*_p_* will have the same sign as 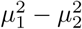. That is, if *μ*_1_ *> μ*_2_, allele *A*_1_ will have a greater advantage over allele *A*_2_ at a maternally expressed imprinted locus than at a paternally expressed one.

The result can perhaps be seen more clearly if we assume that 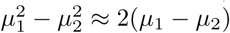, which follows if *μ*_1_ and *μ*_2_ are both close to 1, and we assume that the difference in male and female reproductive variances is the same for both alleles. We then have

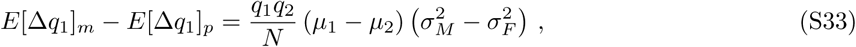

which is our final result discussed below.

### S.5 Two-Generation Model with Imprinting

Here we derive our expression for the expected change in allele frequency *E*[Δ*q*_1_] = *q*_1_*q*_2_(*w*_1_ − *w*_2_), starting from the two-generation effective fitness expressions provided by equation (S31) in the main text and reproduced here:

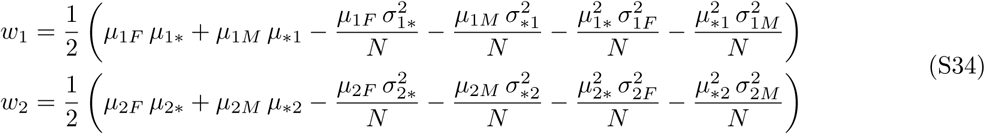

The terms *μ*_1*F*_, *μ*_1*M*_, *μ*_2*F*_, and *μ*_2*M*_ are the mean reproductive success of alleles *A*_1_ and *A*_2_ in females and males, averaged across genotypes. The analogous *σ*^2^ terms are the corresponding reproductive variances.

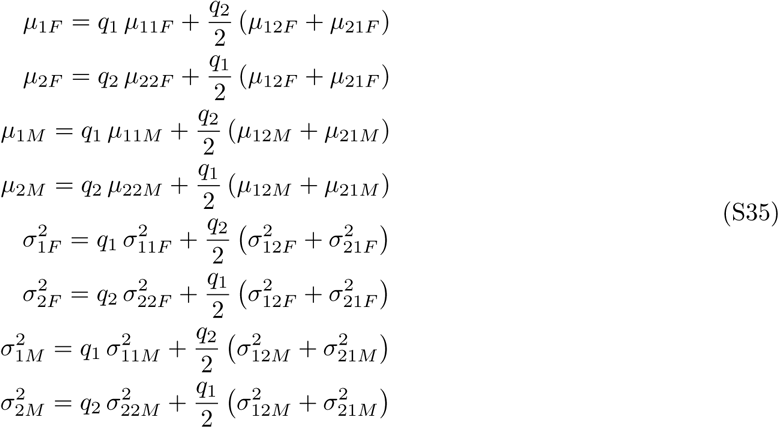

The term *μ*_1*_ represents the mean reproductive success of maternally inherited *A*_1_ alleles, averaged across genotypes and sexes, while *μ*_*1_ is the mean reproductive success of paternally inherited *A*_1_ alleles. The corresponding values for *A*_2_ are given by *μ*_2*\**_ and *μ*_*2_, and again the analogous *σ*^2^ terms are the corresponding reproductive variances.

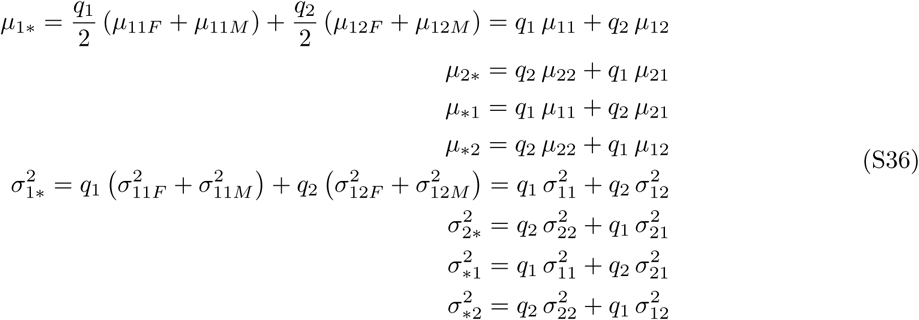

In order to focus our analysis specifically on imprinted gene effects, we will make the simplifying assumption that the alleles do not have sex-specific effects on mean reproductive success (e.g., *μ*11*F* = *μ*11*M* = *μ*11.we now separately consider two cases: an imprinted locus with maternal expression, and an imprinted locus with paternal expression. This allows further simplification. For example, at the maternally expressed locus, *μ*_11_ = *μ*_12_ = *μ*_1_, whereas at the paternally expressed locus, *μ*11 = *μ*21 = *μ*_1_.

Recalling that our values for *μ* were normalized such that the expected mean reproductive output for the population as a whole (and for males and females separately) is one, for the maternally expressed case our expressions for the reproductive means are

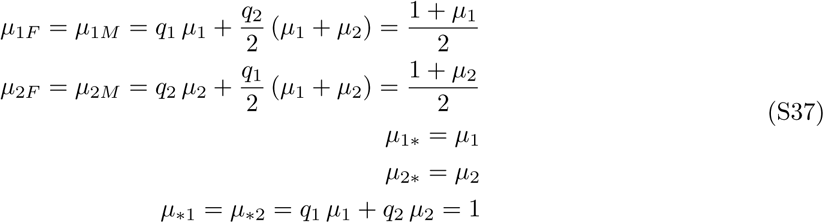

This simplifies our effective fitness expressions to

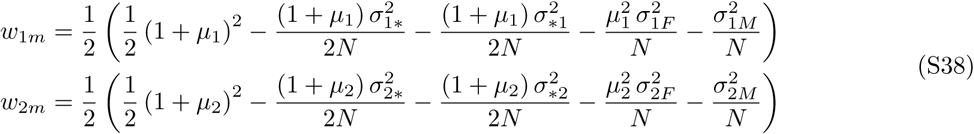

we can also simplify our expressions for reproductive variance. First, we introduce the simplifications associated with assuming that the phenotype depends only on the identity of the maternally inherited allele.

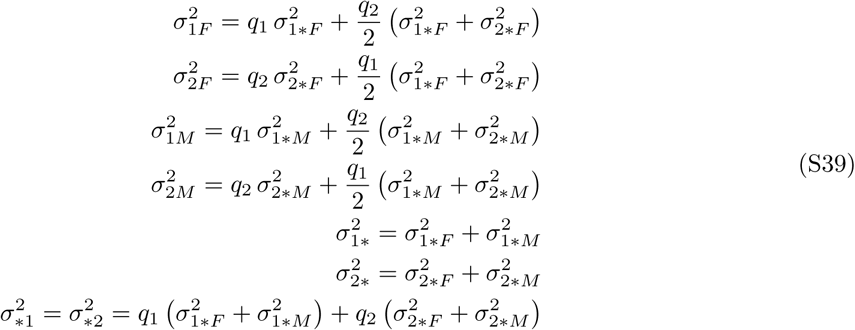

Next, we reparameterize these equations in terms of the total variance (e.g., 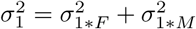) and the difference between male and female variances (e.g.,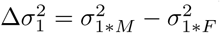). The variance expressions then become

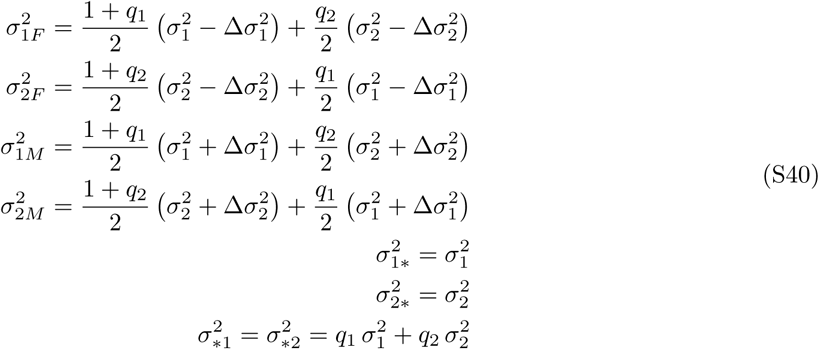

Now, substitution into our fitness expressions gives us

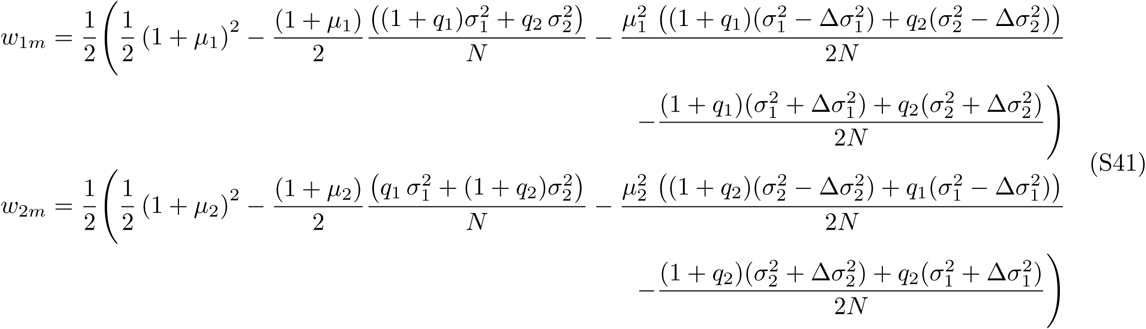

we can contrast these results with the analogous equations for alleles at a paternally expressed imprinted locus. The first terms of the fitness expressions are identical for the two cases. However, the last two terms differ in each case. In the maternally expressed case above, the 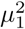 and 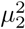 terms are multiplied by the reproductive variances in females. In the paternally expressed case below, these squared mean terms are multiplied by the reproductive variance in males.

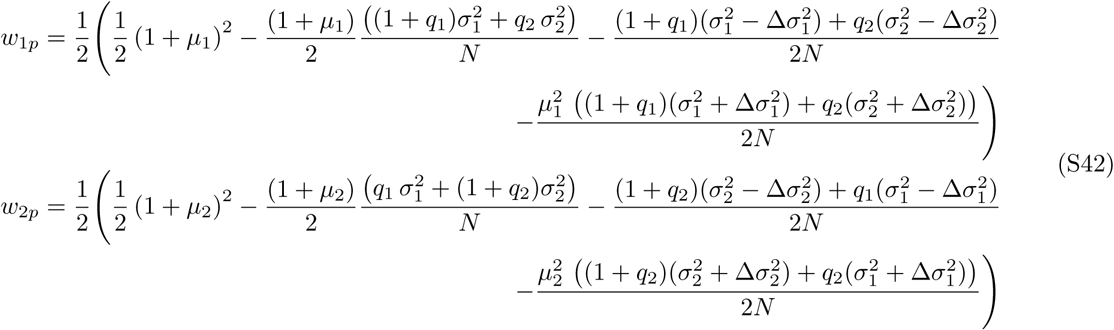

Now, a final substitution will facilitate direct comparison of these results. We set 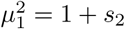 and 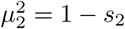. We can then compare the selective advantage of allele *A*_1_ over *A*_2_ at a maternally expressed locus with the advantage of a similar allele at a paternally expressed locus. That is, we consider

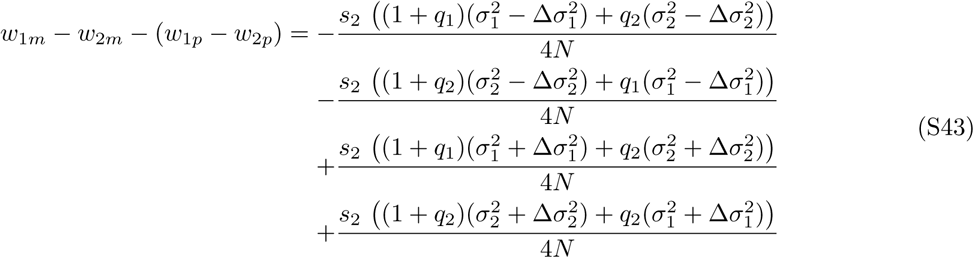

which reduces to

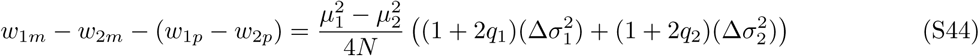

If males have a higher variance of reproductive success than females, as is most often the case, the terms 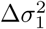 and 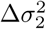 will be positive. That means that, in terms of the relative benefits of increased mean and reduced variance, the benefits of increased mean reproductive success are greater for alleles when they are maternally inherited than when they are paternally inherited.

